# Lipid-mediated reinforcement of FGF/MAPK signaling enables robust otic placode specification

**DOI:** 10.64898/2026.01.06.698011

**Authors:** Stephanie R. Peralta, Natalia Maiorana, Michael L. Piacentino

## Abstract

The formation of cranial placodes requires groups of ectodermal cells to interpret inductive signals in a robust and organized manner, yet how signaling responses are coordinated across a developing field remains incompletely understood. During otic placode specification, fibroblast growth factor (FGF) signaling must overcome intrinsic noise and rising inhibitory feedback to drive collective transcriptional responses, suggesting the existence of reinforcing regulatory events beyond ligand–receptor interactions and gene regulatory networks alone. Here we identify the secreted lipid-binding protein Apolipoprotein D (APOD) as an essential mediator of otic placode specification that links lipid-dependent regulation to FGF/MAPK signaling during early development. We find that *APOD* is strongly expressed in the forming otic placode, where it is necessary for otic specification and morphogenesis. Loss of APOD attenuates ERK1/2 activation, indicating impaired cellular responsiveness to FGF/MAPK signaling. Notably, FGF signaling induces *APOD* expression, establishing a positive feedback loop that reinforces signaling at the tissue level. These findings reveal lipid-mediated regulation of cell signaling as a critical mechanism enabling robust interpretation of developmental signals during sensory placode formation. More broadly, our work highlights lipid management as a key organizational principle by which embryonic tissues achieve coordinated responses to morphogenetic cues.

**Highlights:**

- The lipocalin Apolipoprotein D (APOD) is expressed transiently during early otic development
- APOD is required for specification and morphogenesis of the otic placode
- APOD functions upstream MAPK activation during otic specification
- FGF induces *APOD* expression to build a positive feedback loop

**Graphical Abstract:** 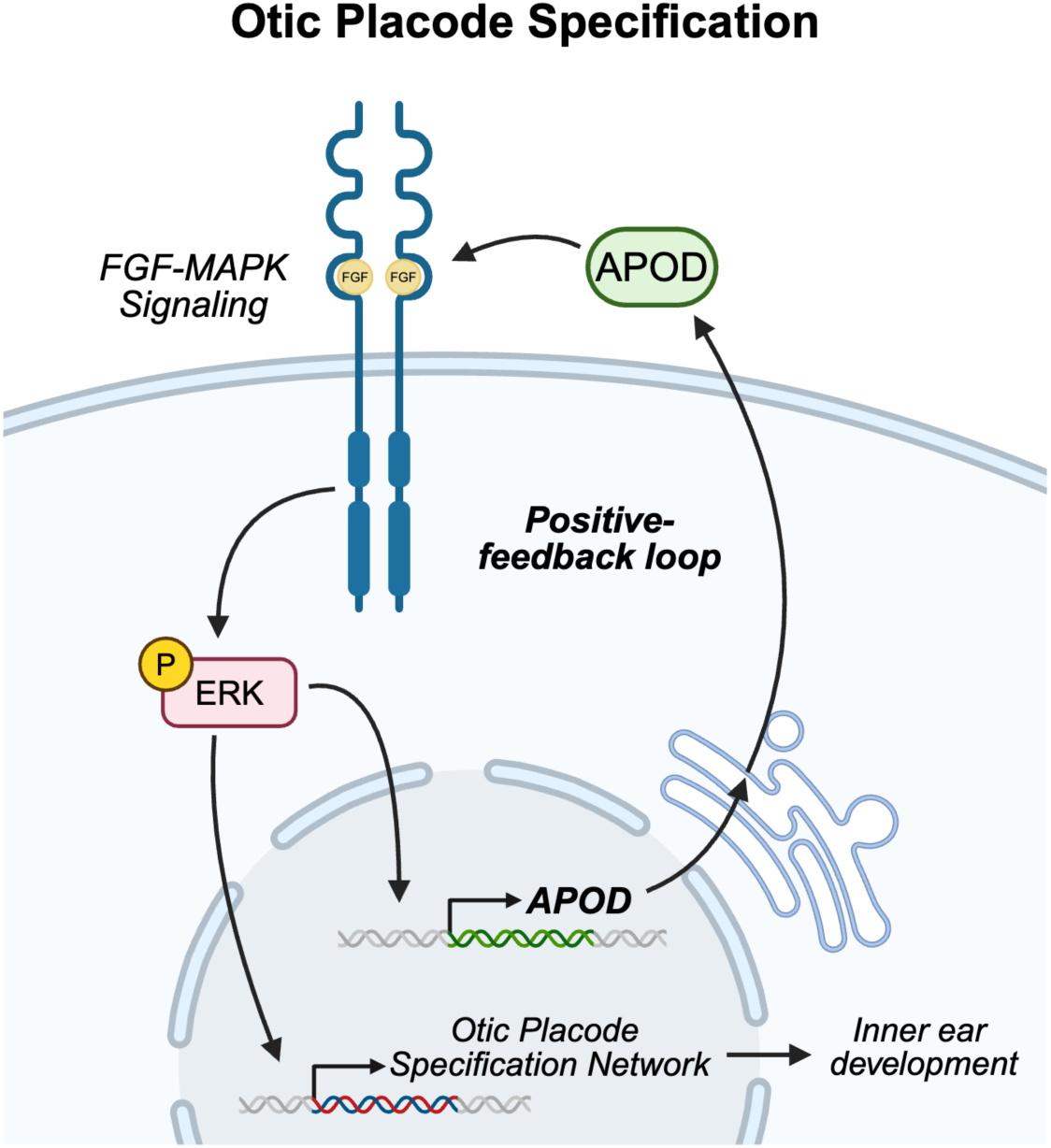

## Introduction

The vertebrate inner ear is an intricate sensory organ responsible for hearing and balance, and arises from a thickened epithelium called the otic placode located adjacent to the developing hindbrain (Baker and Bronner-Fraser, 2001; Sai and Ladher, 2015; Schlosser, 2006; Singh and Groves, 2016; Streit, 2025). During neurulation, the pre-placodal region (PPR) of the anterior neural plate border is subdivided into the distinct neurogenic placodes that give rise to the vertebrate paired sensory organs (Baker and Bronner-Fraser, 2001; Riley, 2021; Singh and Groves, 2016; Streit, 2025). The more posterior otic and epibranchial placodes are established in large part by carefully orchestrated fibroblast growth factor (FGF) signaling. Although much is known about the transcriptional networks and intracellular feedback mechanisms downstream of FGF signaling, less is understood about how groups of ectodermal cells collectively sustain signaling during placode induction. How extracellular factors modulate responsiveness to inductive cues in the face of intrinsic noise and rising inhibitory feedback remains an open question.

After its induction, the PPR becomes subdivided by local signaling interactions that progressively restrict cell fates (Riley, 2021; Sai and Ladher, 2015; Singh and Groves, 2016). First, FGF signals in the posterior region of thePPR induce the bipotent otic-epibranchial precursors (OEPs) characterized by expression of transcription factors including ETS variant transcription factor 4 (*ETV4*, also known as *PEA3*), Paired box gene 2 (*PAX2*), and SRY-box transcription factor 8 (*SOX8*) (Buzzi et al., 2022; Chen et al., 2017; Groves and Bronner-Fraser, 2000; Khatri et al., 2014; Ladher et al., 2000; Lunn et al., 2007; Maroon et al., 2002; Park and Saint-Jeannet, 2008; Streit, 2002; Urness et al., 2010; Wright and Mansour, 2003). Feed-forward circuits that include these factors, together with sustained FGFs and hindbrain-derived Wnt-8c signals, divide the OEPs between medial otic and lateral epibranchial fates (Freter et al., 2008; Ladher et al., 2000; Martin and Groves, 2006; Urness et al., 2010). This results in otic commitment and expression of *SOX10*, which stabilizes otic placode cell survival during complex morphogenetic events including epithelial thickening and invagination (Breuskin et al., 2009; Cheng et al., 2000; Dutton et al., 2009). Finally, the invaginated otic vesicle undergoes further morphogenetic movements to branch and elaborate into the mature vestibulocochlear system and to produce sensory neuroblasts that make the statoacoustic ganglia that relay this sensory information to the brain (Alsina and Whitfield, 2017; Magariños et al., 2012).

FGF signaling is sustained to specify the otic placode through a two-step induction mode, first establishing the initial territory, then recruiting adjacent cells to expand the otic domain (Ladher et al., 2000; Ladher et al., 2005; Martin and Groves, 2006; Riley, 2021; Schimmang, 2007). After secreted FGF ligands bind their transmembrane receptors, they induce phosphorylation through the mitogen activated protein kinase (MAPK) pathway, culminating in activation of the terminal Extracellular Signal-Regulated Kinase 1 and 2 (ERK) (Lunn et al., 2007). Activated ERK potentiates otic development through multiple mechanisms including transcriptional regulation by phosphorylating its targets including ETV4/5 (Garg et al., 2018; Raible and Brand, 2001; Simon et al., 2025), and the AP-1 complex which recruits the p300 histone acetyltransferase to prime otic gene enhancers (Tambalo et al., 2020). FGF activates several members of the otic gene regulatory network through this feed-forward transcriptional regulation (Anwar et al., 2017; Chen et al., 2017; Yang et al., 2013). To prevent overactivation and expansion of FGF responses, these positive feedback mechanisms are counteracted by negative feedback loops. FGFs induce expression of Sprouty members *SPRY1* and *SPRY2* and the dual specificity phosphatase *DUSP3* (also known as *MKP3*), which act intracellularly to inhibit the MAPK cascade (Chambers and Mason, 2000; Ekerot et al., 2008; Lunn et al., 2007; Mahoney Rogers et al., 2011; Urness et al., 2008; Zhang et al., 2014). While these findings illustrate that FGF responses are carefully controlled, we still lack a complete understanding of the extracellular regulatory mechanisms at play during otic development.

Apolipoprotein D (APOD) is a secreted lipocalin with well-established lipid-binding capacity that has been implicated in modulating cell signaling and stress responses in diverse contexts. APOD manages local lipid composition and protects against lipid peroxidation and oxidative stress, thereby influencing lipid metabolism, regulating membrane integrity, and organizing membrane properties (Rassart et al., 2020; Sanchez and Ganfornina, 2021). Importantly, since APOD modulates ERK activation and localization in other contexts (Leung et al., 2004; Sarjeant et al., 2003), APOD may also regulate MAPK responsiveness to FGF signaling during otic placode development.

Here, we show that APOD is required for otic placode specification through positive regulation of FGF/MAPK signaling. Using avian embryos, we find that *APOD* is transiently expressed in multiple craniofacial cell types, most notably the otic placode. Loss-of-function and epistasis experiments indicate that *APOD* is required early in the gene regulatory network underlying otic placode specification and morphogenesis, and that it is necessary to strengthen and sustain MAPK activity throughout induction and specification. Finally, we show that *APOD* itself is a target of FGF signaling. Together our results identify a previously unrecognized extracellular feedback mechanism that sustains FGF/MAPK signaling during otic placode specification and reveals lipid-associated regulation as a key contributor to coordinated interpretation of developmental signals.

## Results

### APOD is transiently expressed in multiple tissues during craniofacial development

We previously identified *APOD* as a bone morphogenetic protein (BMP)-responsive gene expressed in the early migrating neural crest cells in avian embryos (Piacentino et al., 2021). Here we expand our spatiotemporal analysis by examining *APOD* expression using hybridization chain reaction fluorescent in situ hybridization (HCR-FISH) in wild-type chicken embryos. Our analysis revealed little to no expression of *APOD* in the embryonic head at neural plate border stages, but expression was observed in the posterior ingressing mesoderm (Fig. 1A; Hamburger Hamilton (HH) stage 7). We first identified cranial expression of *APOD* in a region adjacent to the future hindbrain at HH8, corresponding to the position of the otic-epibranchial precursor cells (Fig. 1B, OEPs); its expression was then upregulated in the otic placode by HH9 (Fig. 1C). *APOD* expression persisted in the otic placode through HH11 (Fig. 1D,D’’) and was then downregulated by HH13 and 16 (Fig. 1E-F). Interestingly, we also observed more transient expression of *APOD* in additional craniofacial tissues. High *APOD* expression was evident in delaminating cranial neural crest cells, as identified by *SOX10* co-expression, and this expression was retained at low levels in neural crest cells during migration (Fig. 1C,C’). We also found *APOD* expression in a domain of the cranial surface ectoderm (Fig. 1D,D’). Finally, *APOD* expression was detected in the olfactory placode (Fig. 1E), and later in the epibranchial placodes, including presumptive ingressing neuroblasts (Fig. 1F,F’). These expression patterns position *APOD* as a novel potential regulator of early craniofacial development.

**Figure 1.**
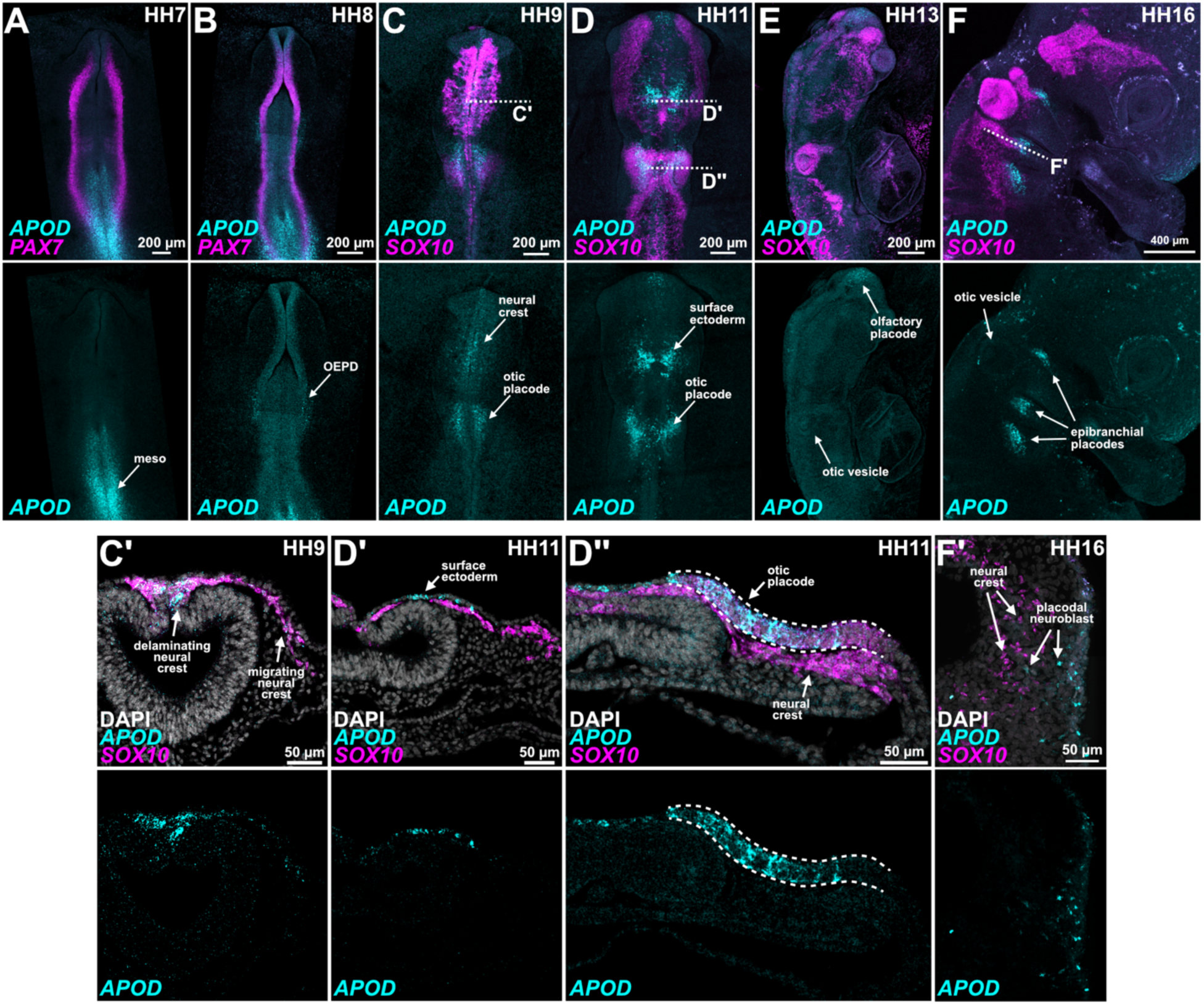
*APOD* is expressed during multiple steps of craniofacial development. Hybridization chain reaction fluorescence *in situ* hybridization probing expression of *APOD* (cyan), neural plate border marker *PAX7* (**A**, **B**; magenta), and neural crest and otic placode marker *SOX10* (**C-F**; magenta) in wild-type chicken embryos at the indicated stages in whole mount (above) and transverse section (below). *APOD* expression was detected in the ingressing mesoderm during gastrulation (**A**, meso), then in the presumptive otic-epibranchial precursors during neurulation (**B**, OEPs). Expression in the otic placode persists through Hamburger Hamilton stage 11 (**C, D, D**’’; HH11), and additional *APOD* is expressed in the medial surface ectoderm over the midbrain (**D, D**’). Transient *APOD* expression was observed in the midbrain neural crest cells, most strongly during delamination but persisting through migration (**C, C’**). By HH13, craniofacial *APOD* expression appears restricted to the rostral olfactory placode, having diminished in expression more caudally (**E**). Finally, *APOD* expression is observed in the epibranchial placodes at HH16 and is maintained as these placodal neuroblasts ingress to interact with migrating neural crest (**F, F’**).

### APOD is required for the formation of the otic placode

Since we saw high expression of *APOD* in the otic placode, we asked if its expression is necessary for otic placode development. We performed unilateral APOD knockdown in chicken embryos by electroporation of an *APOD*-targeting splice-blocking morpholino (MO) on the right side, and a non-targeting control MO on the left side of gastrulation stage embryos (schematized in Fig. 2A). The electroporated embryos were allowed to develop until stage HH10, at which point we performed immunofluorescence to detect expression of SOX10, a transcription factor required for otic placode lineage commitment, survival, and function (Breuskin et al., 2009; Cheng et al., 2000). We observed robust expression of SOX10 in the otic placode domain adjacent to the hindbrain on the control side, indicating appropriate otic placode development at this stage (Fig. 2B, left). Interestingly, we found a dramatic and reproducible loss of SOX10 expression on the contralateral *APOD* knockdown side (Fig. 2B, quantified in Fig. 2C). Importantly, staining for the mitotic marker phospho-histone H3 or the apoptotic marker cleaved caspase 3 revealed no significant difference in proliferation or cell survival in the otic territory (Fig. S1).

**Figure 2.**
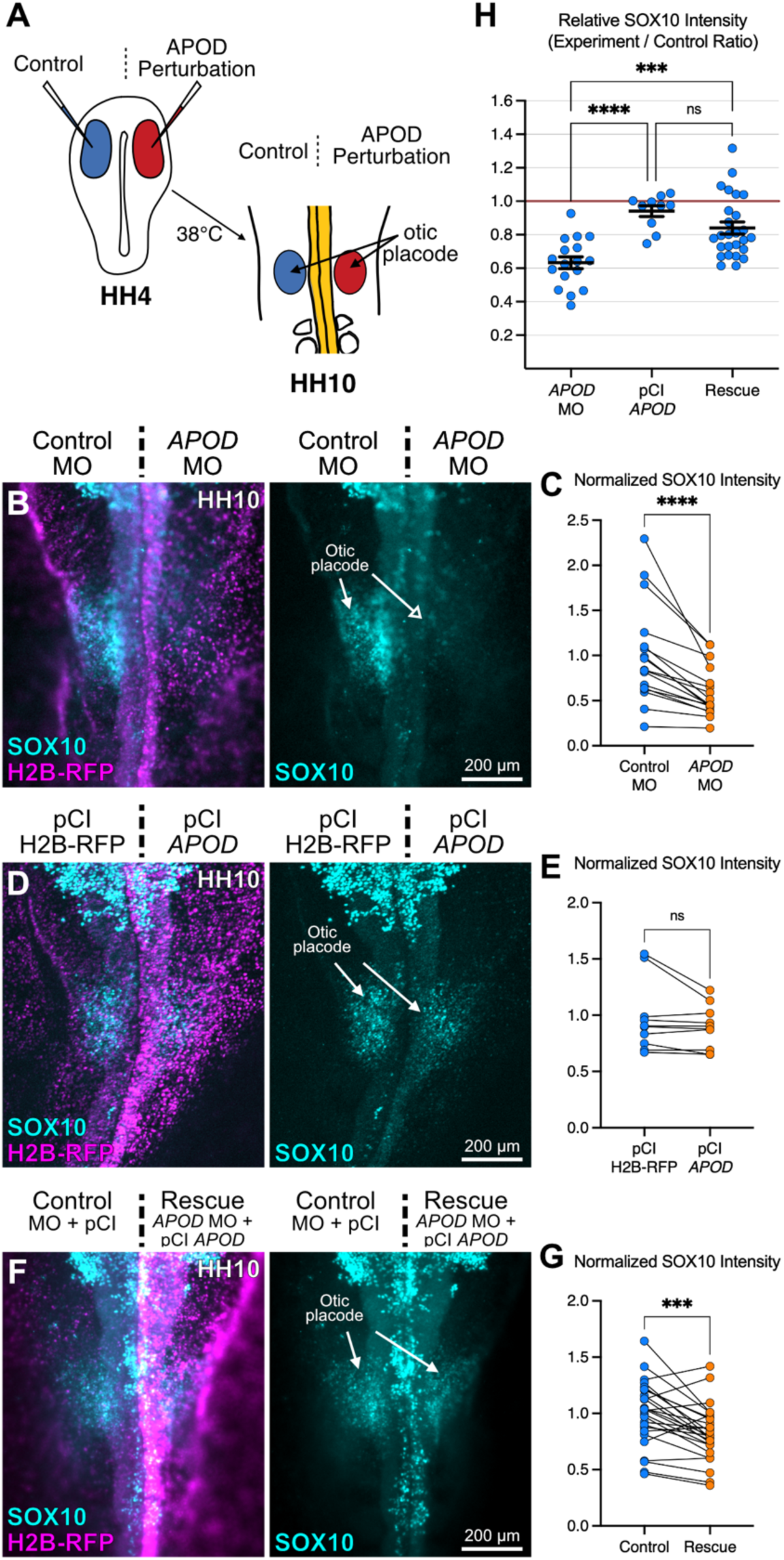
APOD is required for otic placode formation. **A.** Gastrulating chicken embryos were bilaterally electroporated during gastrulation (HH4) with *APOD* perturbation reagents on the right side and the most appropriate control reagents on the contralateral left side. Embryos were grown to HH10 at stained for SOX10 using immunofluorescence to assess otic placode formation. **B, D, F**. Representative dorsal view images of HH10 embryos after the indicated treatments, displaying H2B-RFP co-electroporation confirms successful transfection. **C, E, G**. Slope charts displaying normalized mean SOX10 immunolabeling paired between the embryo sides. Statistical annotations reflect Wilcoxon matched pairs signed rank tests. **H**. Ratio of experimental over control SOX10 intensity measurements from B, E, and G. Statistical annotations reflect Tukey’s multiple comparisons testing after one-way ANOVA. **** *P*<0.0001, *** *P*<0.001, ns not significant.

We next asked if APOD is sufficient to promote otic fates in the cranial ectoderm. We cloned the full-length chicken *APOD* coding sequence into an overexpression vector harboring the ubiquitous CAG promoter to overexpress APOD alongside an H2B-RFP lineage tracer (pCI *APOD*). We examined the effects of APOD overexpression on SOX10 immunoreactivity in the otic placode and compared this to the contralateral embryo side, carrying the same plasmid but lacking the *APOD* coding sequence (pCI H2B-RFP). The results showed that APOD overexpression did not have a detectable impact on SOX10 expression in the otic placode (Fig. 2D, E). Since the pCI *APOD* construct is insensitive to the splice-blocking *APOD* MO, we then co-electroporated these reagents together to test their specificity (Fig. 2F, G). To compare the magnitude of these effects between manipulations, we analyzed experimental/control SOX10 intensity ratios (Fig. 2H). This analysis revealed a partial but significant rescue of SOX10 expression in co-electroporated embryos compared to *APOD* MO alone, suggesting that the observed knockdown effects are specific to loss of APOD and not off-target effects. Together, our results demonstrate that *APOD* function is required for proper otic placode formation.

### Otic vesicle formation requires APOD function

To distinguish if the effect of *APOD* loss reflects reduced expression of a single gene, or a more significant disruption to otic placode development, we next examined transverse sections of *APOD* morphants at the level of the otic placode. At this stage, the placode ectoderm thickens into a more columnar epithelium before invaginating to form the otic vesicle (Anniko and Wikström, 1984; Bancroft and Bellairs, 1977; Christophorou et al., 2010). Transverse sections confirmed a reduction in SOX10-positive cells after APOD loss but also displayed atypical placode morphogenesis (Fig. 3A). While the otic placode on the control side exhibited the typical columnar epithelial morphology, the otic placode after *APOD* knockdown instead appeared more cuboidal, reminiscent of naïve cranial ectoderm.

**Figure 3.**
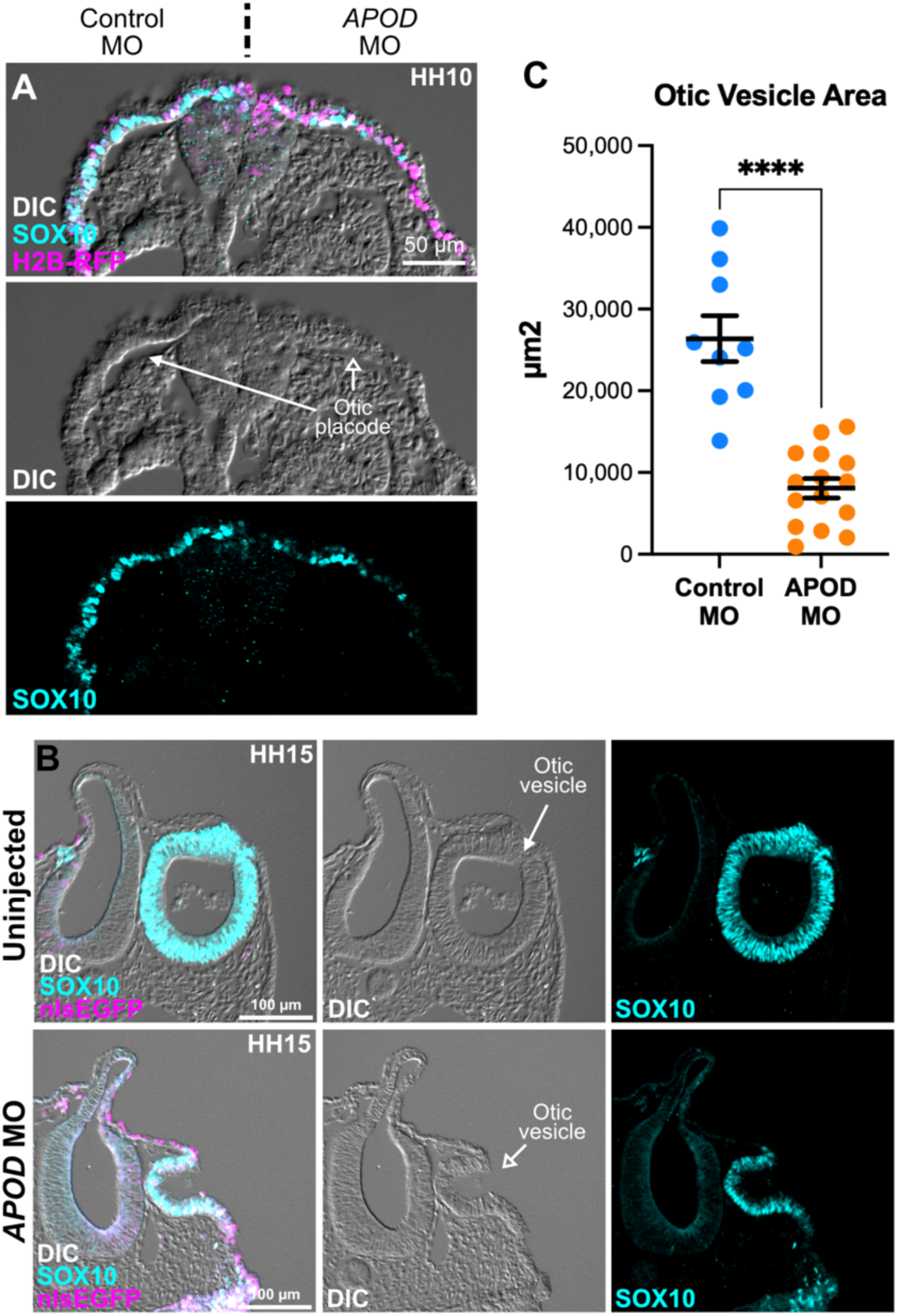
APOD is necessary for appropriate otic vesicle development. **A**. Transverse section through the otic placode of an APOD morphant embryo at stage HH10. Differential interference contrast (DIC) imaging reveals appropriate columnar morphology of the otic placode on the left, control side, with more disorganized and squamous epithelial morphology on the right (closed arrow), APOD-deficient side (open arrow). **B.** Transverse sections through the otic vesicle on the uninjected (top) and *APOD* MO injected (bottom) sides of the same embryo at stage HH15 illustrates reduced tissue area, delayed fission from the epidermis, and reduced SOX10 expression. **C**. Quantification of otic vesicle cross sectional area in Control MO and *APOD* MO transfected vesicles. ** *P*<0.01, Welch’s unpaired t-test.

This result could reflect either a developmental delay or a critical requirement for *APOD* function. To distinguish between these possibilities, we performed unilateral electroporation of Control or *APOD* MOs and examined the resulting otic vesicle size after they have largely invaginated at stage HH13-HH14 (Bancroft and Bellairs, 1977). After immunostaining for SOX10, *APOD* morphant otic vesicles in transverse section were significantly smaller than those from stage-matched control embryos and contralateral uninjected sides (Fig. 3B). Using confocal microscopy in whole mount embryos, we found a significant and reproducible reduction in otic vesicle cross-sectional area after APOD knockdown (Fig. 3C). Interestingly, at these later stages we did not detect expression of *APOD* mRNA (Fig. 1E,F), suggesting that this phenotype reflects an earlier requirement for APOD. These results strongly implicate APOD in the formation of the otic placode.

### APOD is required for otic placode specification

To decipher which step in development *APOD* feeds into *SOX10* expression, we examined the upstream gene regulatory network underlying otic placode induction and specification. Previous network analyses have defined the expression of transcription factors including *ETV4*, *PAX2*, and *SOX8* as expressed during the induction of the OEPs, then directly regulate *SOX10* expression during otic commitment (Betancur et al., 2011; Chen et al., 2017; Groves and Bronner-Fraser, 2000; Lunn et al., 2007; McKeown et al., 2005; Streit, 2002). We examined the spatial expression pattern of these factors, as well as *FOXI3* which is initially expressed in the OEPs, then transitions to the more lateral epibranchial and more anterior trigeminal placode territory (Khatri and Groves, 2013; Urness et al., 2010). We confirmed the expression of these transcription factors in the otic placode and adjacent tissues using HCR-FISH in wild-type chicken embryos at HH10 (Fig. 4A).

**Figure 4.**
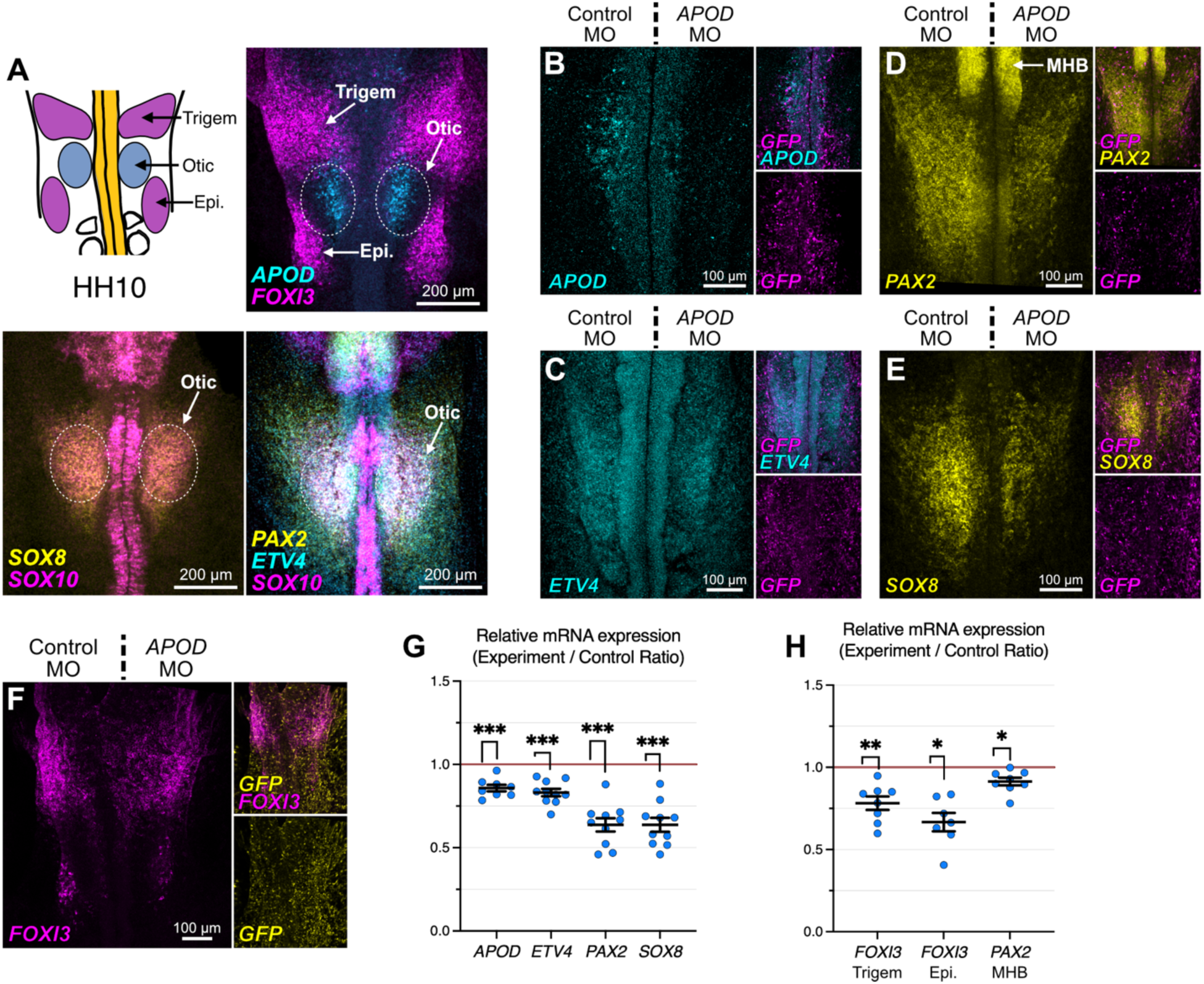
Otic placode specification networks require APOD. **A.** Cartoon schematic and images of HH10 wild-type chicken embryos processed by HCR-FISH for *APOD* expression alongside otic placode expression of *SOX8, PAX2, ETV4,* and *SOX10*, and expression of *FOXI3* in the epibranchial (Epi.) and trigeminal (Trigem) placodes. **B-F.** Representative images showing HCR-FISH results for the indicated markers after right side *APOD* MO compared to left side Control MO electroporations. *GFP* expression was included to assess successful transfection. MHB, midbrain-hindbrain boundary. **G-H.** Relative mRNA expression calculated as experimental/control side intensity for the indicated genes within the otic placode domain (G), and in the adjacent tissues (H). *** *P*<0.001, ** *P*<0.01, * *P*<0.05, Paired t-test.

By measuring the expression of these genes after APOD knockdown using HCR-FISH and comparing to the contralateral control, we found reduced expression of *APOD, ETV4, PAX2,* and *SOX8* in the otic placode (Fig. 4B-E, 4G). Further, Reduced *APOD* mRNA levels illustrate the efficiency of the splice-blocking *APOD* MO. We also observed significant reduction in *FOXI3* expression both in the nearby epibranchial and trigeminal placodes, and reduced *PAX2* expression in the midbrain-hindbrain boundary (MHB) (Fig. 4D,F,H), suggesting that *APOD* has non-cell autonomous effects that may result from its role as a secreted lipid transporter. Since *ETV4, PAX2*, and *SOX8* act upstream *SOX10* expression, and *APOD* is coincidently expressed at HH8 (Fig. 1B, Fig. 6A, see also (Betancur et al., 2011; Chen et al., 2017; McKeown et al., 2005)), these results suggest that *APOD* acts early in the onset of otic placode specification, and its effects on SOX10 during otic fate commitment at HH10 is likely a result of earlier function.

### Otic specification subcircuits partially rescue otic specification

We next explored if any of these otic specifier genes are sufficient to rescue SOX10 expression in the absence of APOD. We performed epistasis analyses by co-electroporating *APOD* MO with plasmids driving constitutive expression of *ETV4*, *PAX2*, or *SOX8*. Electroporated embryos were then immunostained for SOX10 to assess otic placode specification at HH10 (Fig. 5A-C). Compared with *APOD* MO alone, we did not observe a significant increase in SOX10 immunoreactivity by misexpression of *ETV4*, *PAX2*, or *SOX8* (Fig. 5D). Interestingly, we did note a trend toward even further reduced SOX10 expression in PAX2-overexpressing embryos, consistent with its auto-inhibitory effects and insufficiency to confer otic fates (Chen et al., 2017; Christophorou et al., 2010). We also observed a slight, albeit not significant increase in SOX10 expression after overexpression of SOX8. In these experiments we found SOX10 ectopically expressed throughout the ectoderm, consistent with the recently reported role for SOX8 in remodeling cranial ectoderm toward an otic fate (Buzzi et al., 2022).

**Figure 5.**
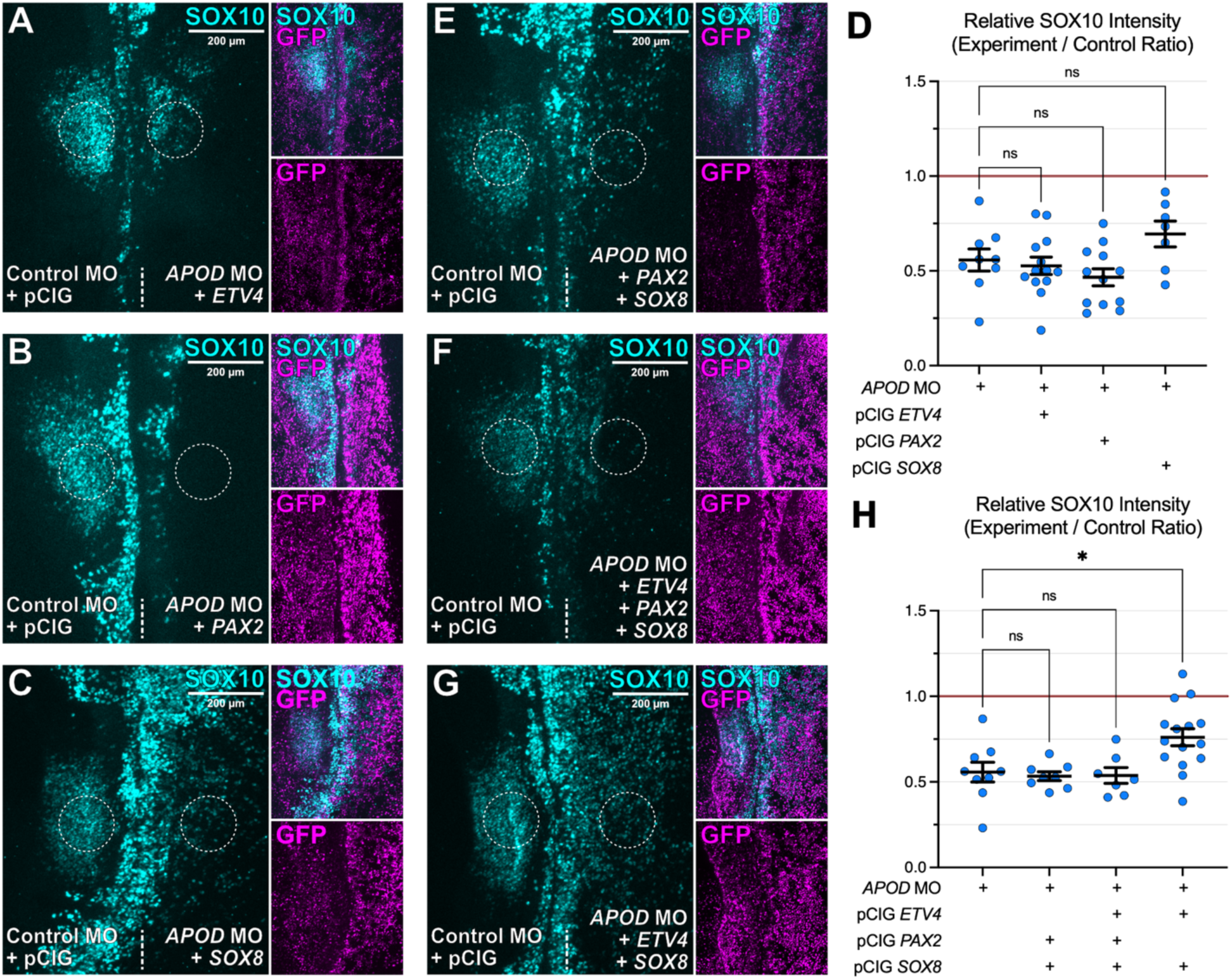
An otic subcircuit partially rescues the SOX10 expression observed with APOD loss. A-C, E-F. Representative images from epistasis experiments in which *APOD* MO was co-electroporated with DNA constructs to overexpress the indicated transcription factors (alone (A-C) and in combination (E-G)) and assess sufficiency to rescue SOX10 expression. **D, H**. Relative SOX10 immunofluorescence intensity displayed as the ratio of experimental/control side intensities. * *P*<0.05, ns not significant, One-way ANOVA with Tukey’s HSD comparisons against *APOD* MO alone.

Other efforts to reprogram cell fates during similar developmental processes have required a subcircuit of transcription factor (Gandhi et al., 2020; Simoes-Costa and Bronner, 2016); this led us to ask if combinations of *ETV4*, *PAX2*, and/or *SOX8* expression instead can rescue SOX10 expression. We repeated these experiments with various combinations and observed a significant but partial restoration of SOX10 expression levels in the otic domain when driving *ETV4* and *SOX8* expression (Fig. 5E-H). This observation further strengthens our interpretation that the observed SOX10 expression could be due to the reprogramming effect of SOX8.

### APOD is required for MAPK pathway activation

Induction, specification, and patterning of the otic placode is critically dependent on the FGF/MAPK cascade (Hans et al., 2007; Lunn et al., 2007; Martin and Groves, 2006; Urness et al., 2010). Interestingly, APOD has been shown in other contexts to regulate ERK activation or localization in response to growth factor signaling, thereby impacting pathway function (Leung et al., 2004; Sarjeant et al., 2003); this prompted us to ask if APOD acts to regulate MAPK signaling in the otic placode. To test this hypothesis, we analyzed the dual phosphorylation and activation of ERK using phospho-specific immunofluorescence (dpERK1/2, Fig. 6A-D). The results show reduced MAPK activity across multiple stages of otic placode development as early as induction of the OEPs (5-6ss), specification (7-8ss), and commitment (9-10ss).

**Figure 6.**
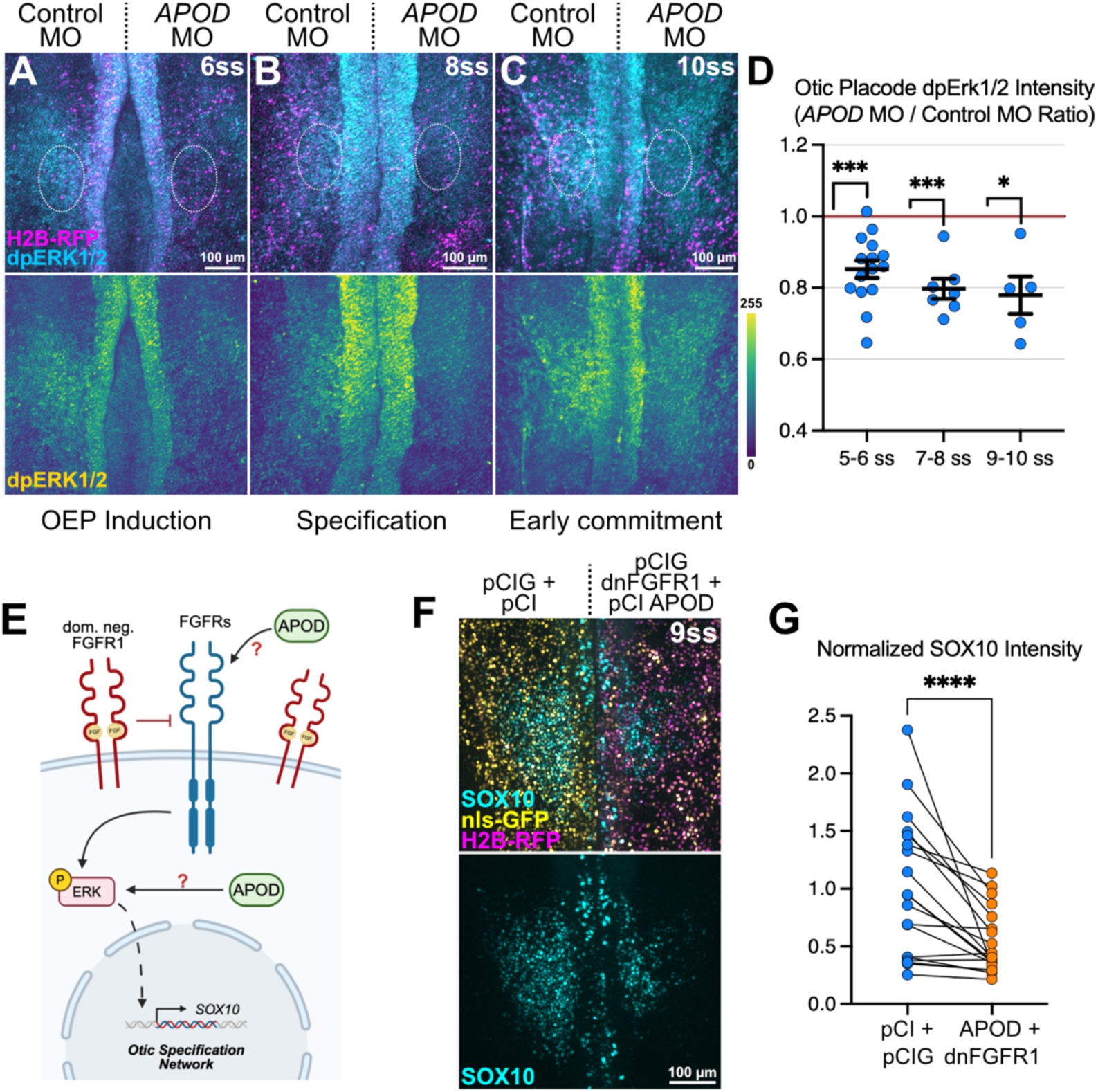
APOD is required for appropriate MAPK activation during otic placode specification. A-C. Representative images of *APOD* morphant embryos at 6, 8, and 10 somite stages after immunofluorescence for the activated, dually phosphorylated ERK1/2 MAP kinase (dpERK1/2, bottom panels pseudocolored according to the Viridis scale on right). **D.** Relative ERK1/2 phosphorylation displayed as the ratio of experimental/control side intensity within the otic placode domain at the indicated stages. *** *P*<0.001, * *P*<0.05, Paired t-test. **E.** Schematic illustration of FGFR-dependent phosphorylation of ERK1/2 and subsequent otic placode specification. Truncated dnFGFR1 sequesters FGF ligands and preventing endogenous FGFR activation. **F.** Misexpression of dominant negative FGF receptor 1 (dnFGFR1) together with APOD shows loss of SOX10 expression in the otic territory. **G.** Slope charts displaying normalized mean SOX10 immunolabeling paired between the embryo sides. **** *P*<0.0001, Wilcoxon matched pairs signed rank test.

Since APOD function was required for MAPK activation in the otic placode, we next sought to ask where in the FGF/MAPK cascade APOD functions. We blocked FGF signaling using a truncated dominant negative FGF receptor (dnFGFR1, (Stuhlmiller and García-Castro, 2012)) which we co-expressed with ectopic APOD. We reasoned that if APOD functions intracellularly between the receptor and phosphorylation of ERK, it would restore MAPK function and subsequent SOX10 expression. Alternatively, if APOD interacts with the FGF ligand or receptor extracellularly, the dnFGFR1 effect would be dominant (Fig. 6E). The results showed SOX10 loss on the experimental side, compared to contralateral controls (Fig. 6F-G), supporting a model in which APOD positively regulates MAPK signaling at, or above, the level of FGF receptor activation (Fig. 6E), consistent with the reported roles of secreted APOD in regulating MAPK pathways (Leung et al., 2004; Sarjeant et al., 2003). Thus, APOD represents a novel positive regulator of MAPK signaling in the otic placode, necessary for specification of this tissue.

### FGF signaling initiates APOD expression completing a positive feedback loop

By examining published bulk RNA-sequencing data, we observe *APOD* expression coincides with the upregulation of *FGF8* and *PAX2* expression, then closely follows expression dynamics of *FGF8* and its targets *ETV4* and *MKP3* (Fig. 7A, data from (Chen et al., 2017)). This raised the possibility that *APOD* itself is an FGF target. FGF/MAPK signaling phosphorylates the AP-1 complex to localize p300 and deposit H3K27ac marks on OEP enhancer sequences, thereby opening local chromatin and priming cells for otic fates (Tambalo et al., 2020). Thus, we examined these FGF-responsive otic enhancer peaks and interestingly found the proximal promoter of *APOD* shows a peak in local H3K27ac marks in the FGF2-treated samples that was absent from naïve PPR ectoderm (Fig. 7B, data from (Tambalo et al., 2020)). This is consistent with other reports showing that *APOD* contains ERK1/2-sensitive SRE-1 and AP-1 sites in its proximal promoter (Do Carmo et al., 2007; Sanchez and Ganfornina, 2021), and suggests that *APOD* expression in the otic placode may activated by FGF-dependent AP-1 function.

**Figure 7.**
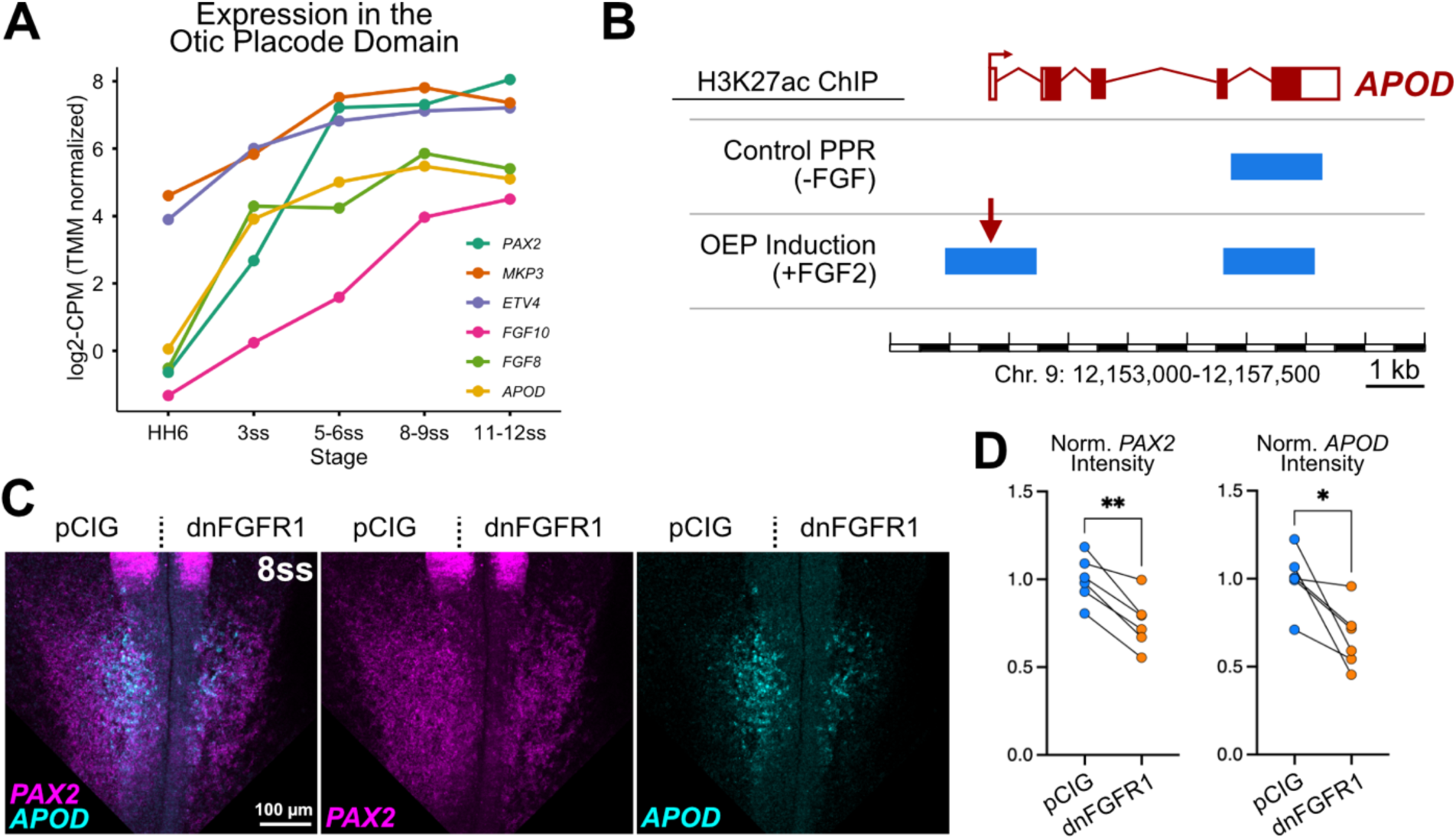
*APOD* expression requires FGF activity. **A.** Bulk RNA sequencing from dissected otic placodes shows *APOD* expression follows similar trends to the *FGF8* ligand, and its targets *ETV4* and *MKP3*. Raw data from (Chen et al., 2017). **B.** Genome browser indicating positions of FGF-responsive putative otic gene enhancers as determined by differential H3K27ac ChIP-seq in dissected naïve preplacodal ectoderm (PPR) and PPR explants induced toward otic-epibranchial precursor identity by addition of FGF2. Raw data from (Tambalo et al., 2020). The results show a significant peak in H3K27 acetylation at the *APOD* promoter compared to naïve controls (red arrow). **C.** Representative images of dnFGFR1-electroporated embryos at 8ss processed by HCR-FISH to show expression of *PAX2* and *APOD* mRNAs. **D**. Slope charts displaying normalized mean *PAX2* and *APOD* mRNA expression in the otic territory at 8-9ss, paired between embryo sides. ** *P*<0.01, * *P*<0.05, Paired t-test.

To further test this hypothesis, we inhibited FGF/MAPK signaling by expressing the dnFGFR1 and performed HCR-FISH to assay expression of *PAX2* and *APOD*. Experiments reveal significant downregulation of both transcripts in the prospective otic territory when FGF signaling is attenuated (Fig. 7C,D), confirming that FGF/MAPK signaling acts upstream *APOD* expression. Together, our results indicate that *APOD* expression is induced by FGF/MAPK signaling in the otic placode, then acts in an extracellular positive feedback loop to strengthen MAPK activation in the cranial ectoderm. This positive feedback loop is essential for appropriate specification and commitment of otic placode fates during early ear development.

## Discussion

Precise tuning of FGF/MAPK signaling is essential for otic placode induction and specification (Ekerot et al., 2008; Freter et al., 2008; Ladher et al., 2000; Mahoney Rogers et al., 2011; Martin and Groves, 2006; Riley, 2021; Urness et al., 2008; Yang et al., 2013), yet extracellular mechanisms that sustain signaling against rising negative feedback remain poorly understood. Here we identify APOD, a secreted lipocalin, as a novel extracellular positive regulator that reinforces MAPK activity and is required for otic placode specification in avian embryos. We find that *APOD* is transiently expressed in the otic placode and neighboring craniofacial tissues, its expression requires FGF/MAPK activity, and loss of APOD reduces ERK output and otic gene expression during induction and specification. Together these results define an extracellular positive feedback loop that complements known intracellular and transcriptional feedback circuits, revealing an additional layer of regulation that supports robust interpretation of FGF signals during placode development.

FGF signaling drives both positive and negative feedback mechanisms to regulate its activity. Feed-forward transcriptional programs act through FGF/MAPK-dependent epigenetic priming and induction of self-reinforcing otic gene regulatory networks, thereby amplifying and stabilizing otic identity (Anwar et al., 2017; Chen et al., 2017; Tambalo et al., 2020; Yang et al., 2013). At the same time, FGF-induced *SPRY1/2* and *DUSP6* act as intracellular negative regulators to attenuate MAPK output (Ekerot et al., 2008; Lunn et al., 2007; Mahoney Rogers et al., 2011; Urness et al., 2008). Our data place APOD within this regulatory network as an extracellular component of FGF-driven positive feedback. We find that FGF activates expression of *APOD* (Fig. 7), consistent with past evidence of AP-1-dependent activation of *APOD* expression in response to lipopolysaccharide stimulation (Do Carmo et al., 2007). This relationship suggests that *APOD* expression is both a consequence and potentiator of MAPK activity. Accordingly, while reduced APOD transcript detection following splice-blocking morpholino treatment (Fig. 4B,G) may reflect nonsense-mediated decay of mis-spliced transcripts, it could also arise indirectly from diminished MAPK activation and reduced *APOD* transcription. Because APOD is secreted (Leung et al., 2004; Rassart et al., 2020; Sarjeant et al., 2003), this extracellular feedback loop may act in a paracrine manner to help sustain and potentially propagate ERK activity across the developing placodal field as as intracellular inhibitors accumulate.

Our epistasis experiments showed that APOD failed to rescue dominant negative FGFR1, suggesting that APOD acts at or above the level of receptor activation during otic development (Fig. 6E-G). These data suggest a model in which secreted APOD functions in the extracellular environment to potentiate FGF signaling competence, rather than acting downstream of MAPK activation. FGF ligands are often bound, stabilized, and delivered to their receptors by cell surface heparin sulfate proteoglycans (HSPGs)(Aviezer et al., 1994; Duchesne et al., 2012; Venero Galanternik et al., 2015). The lipocalin family members β-lactoglobulin and Fel d 4 are internalized in an HSPG-dependent manner (Habeler et al., 2020), raising the possibility that APOD may engage HSPG-associated signaling complexes at the cell surface. Alternatively, APOD is internalized by the ubiquitously expressed transmembrane receptor Basigin (BSG, also known as CD147)(Iacono et al., 2007; Najyb et al., 2015). Since BSG can also activate the MAPK pathway (Li et al., 2023), we cannot exclude a model in which APOD acts in parallel to FGFs, with both pathways converging on ERK activation. Additional biochemical studies will be needed to clarify this mechanism and identify the APOD binding partners in the context of otic development.

Through a series of epistasis experiments, we found that overexpression of otic specifier genes *ETV4, PAX2*, or *SOX8* alone or together was insufficient to restore otic placode development in the absence of APOD (Fig. 5). We did observe ectopic SOX10 expression when *SOX8* was misexpressed, consistent with recent reports that show *SOX8* is sufficient to reprogram the cranial ectoderm toward an otic identity (Buzzi et al., 2022). However, this does not reflect a true rescue as SOX10 was scattered throughout the cranial ectoderm and did not organize into an appropriately patterned ectodermal placode. These findings suggest that APOD is required not simply for induction of otic transcriptional programs, but for establishing the signaling context necessary for the coordinated spatial deployment. This raises the possibility that APOD regulates otic placode development through one or more pathways parallel to, or upstream of, the FGF-dependent MAPK cascade. For example, in other contexts, APOD regulates cell growth and is upregulated upon growth arrest (Do Carmo et al., 2007; Rassart et al., 2020; Sarjeant et al., 2003). However, the absence of detectable changes in proliferation or apoptosis after APOD knockdown (Fig. S1) argues that APOD-dependent MAPK activity in the otic placode primarily governs cellular identity rather than tissue growth.

Our results also suggest additional roles for APOD in craniofacial development. *APOD* is expressed in the cranial neural crest, olfactory and epibranchial placodes, and gastrulating mesoderm (Fig. 1), all of which undergo complex morphogenetic movements such as ingression, migration, and invagination (Gao et al., 2025; Piacentino et al., 2020; Schlosser, 2006; Zhang et al., 2025), raising the possibility of conserved molecular functions across these contexts. Consistent with this, *APOD* expression has been reported in the developing mouse cranial mesenchyme surrounding the neural tube and olfactory epithelium, potentially reflecting otic vesicle-derived neuroblasts that contribute to the statoacoustic ganglion (Sánchez et al., 2002). Later, *APOD* is expressed in the mature cochlea, and *APOD* knockout mice show a trend toward reduced hearing acuity at certain frequences, although these differences did not reach statistical significance (Hildebrand et al., 2005). No overt morphogenetic defects have been reported, possibly due to genetic compensation by other lipocalins, but these observations support a conserved role for APOD in otic development and function. In our avian system, loss of APOD reduces *FOXI3* and *PAX2* expression in neighboring placodal and neural territories—where *APOD* transcripts are not detected—respectively (Fig. 4H), suggesting that secreted APOD acts non-autonomously to coordinate signaling states across craniofacial tissues and synchronize their development.

In conclusion, our results describe a novel role for the lipocalin APOD in regulating early inner ear formation, where it functions in a positive feedback loop with FGF/MAPK signaling. By binding and managing membrane lipids, we speculate that APOD may regulate FGF/MAPK signaling through one or more mechanisms. For example, APOD may contribute to partitioning lipid microdomains on the plasma membrane that facilitate ligand-receptor engagement, sequester arachidonic acid or other lipid modulators to tune feedback regulation, or by internalization and recruitment to lysosomal membranes, APOD may even influence FGF receptor stability and trafficking (Sanchez and Ganfornina, 2021). Regardless of mechanism, these findings establish APOD as a critical extracellular regulator that ensures robust, coordinated otic specification, and highlight secreted lipocalins as underappreciated organizers of signaling responses during craniofacial morphogenesis.

## Acknowledgments

We would like to thank Noah Gurley for his contributions to application of the phospho-Erk1/2 immunostaining protocol and initiating this line of experimentation, and Dr. Deborah Andrew for helpful comments. This work was funded by the National Institute for Dental and Craniofacial Research award R00 DE029240 (MLP) and Diversity Research Supplement (SRP), and startup funding from the Johns Hopkins University School of Medicine (MLP). ChatGPT (OpenAI) was used to generate initial drafts of data analysis code and for assistance in organizing the structure of the Introduction and Discussion. All code, analyses, and text were reviewed, validated, and finalized by the authors.

## Code and Data Availability

All study data are included in the main text and/or supporting information, with analysis codes and associated source data available on GitHub: https://github.com/PiacentinoLab/2025_Peralta_APOD_Otic/.

## Author Contributions

Conceptualization: S.R.P. and M.L.P.

Experiment design: S.R.P. and M.L.P.

Experimentation: S.R.P., N.M., and M.L.P.

Data analysis: S.R.P., N.M., and M.L.P.

Data interpretation: S.R.P., N.M., and M.L.P.

Manuscript preparation: S.R.P. and M.L.P.

Manuscript editing: M.L.P.

## Materials and Methods

### Embryos and perturbations

Fertilized white leghorn chicken eggs were obtained from commercial sources (University of Connecticut and Michigan State University). *Ex ovo* electroporations were performed at gastrula stage (HH4) using five pulses of 5.8 V for 50 ms at 100 ms intervals. Embryos were then cultured in albumin with 1% penicillin/streptomycin and incubated at 37.8°C to the desired HH stage (Williams and Sauka-Spengler, 2021). Embryos grown to stages HH15-16 were unilaterally electroporated and cultured on agar pads composed of 0.4% BactoAgar, 15.4 mM NaCl, 0.0015% D-Glucose, and 0.1% penicillin/streptomycin in 50% albumin. For transverse sections, fixed embryos were incubated in 5% sucrose for 15 min at room temperature, 15% sucrose overnight at 4°C, 7.5% gelatin at 39°C overnight, then flash-frozen in liquid nitrogen and stored at -80°C. Embryos were then cryosectioned at a thickness of 16 µm. All reagents electroporated either expressed a fluorescent protein or were co-electroporated with nuclear GFP- or RFP-expression constructs (Betancur et al., 2011; Megason and McMahon, 2002); fluorescent protein expression was used to screen for electroporation efficiency and only embryos with high efficiency were included in the analysis. Morpholinos (MOs) were electroporated at 1.0 mM and synthesized by Gene Tools, LLC with the following sequences: control MO: CCTCTTACCTCAGTTACAATTTATA; splice-blocking *APOD* MO: CACAGCGGCACTTACAGCATCTCCT. pCI::*APOD*-ires-H2B-RFP, pCIG::*ETV4*, pCIG::*SOX8*, pCIG::*PAX2*, and pCIG::*dnFGFR1* (Stuhlmiller and García-Castro, 2012) constructs were electroporated individually at 2.0µg/µl. For epistasis experiments, injection mixes were assembled containing 1 µg/µl of each plasmid.

### Hybridization Chain Reaction Fluorescence In Situ Hybridization

Reagents for third-generation hybridization chain reaction (HCR) probes and supplies were purchased from Molecular Instruments or as IDT oPools, and hybridization experiments were performed as previously described (Piacentino et al., 2022). Probe sets were designed against *APOD* (B5 initiator), *SOX10* (B3 initiator), *PAX2* (B2 initiator), *ETV4* (B4 initiator), *SOX8* (B4 initiator), and *FOXI3* (B2 initiator) and detected using corresponding amplifier hairpins labeled with Alexa Fluor 488, Alexa Fluor 546, and Alexa Fluor 647.

### Immunohistochemistry

To perform immunofluorescence analysis, embryos were fixed for 15 min at room temperature in 4% PFA in phosphate buffer, and all subsequent washes and incubations were performed in TBST + Ca^2+^ (50 mM Tris HCl, 150 mM NaCl, and 1 mM CaCl_2_, 0.5% Triton X-100). Blocking was performed in 10% donkey serum for 1 hour at room temperature, and primary and secondary antibody incubations were carried out in 10% donkey serum for one night at 4°C, except for rabbit Phospho-p44/42 MAPK (Erk1/2)(Thr202/Tyr204), for which both primary and secondary incubations required three nights each. Primary antibodies used in this study include rabbit IgG anti-Sox10 (1:500, Millipore Sigma #HPA068898), rabbit Phospho-p44/42 MAPK (Erk1/2)(Thr202/Tyr204) (1:50; Cell Signaling #9101), rabbit anti-phosphohistone H3 (1:250; Millipore #06-570), rabbit anti-cleaved caspase 3 (1:250; R&D Systems #AF835), goat anti-GFP (1:500; Rockland #600-101-215M), and rabbit anti-RFP (1:500; MBL #PM005), and species-specific AlexaFluor-conjugated secondary antibodies raised in donkey (Invitrogen) were used to detect primary antibodies.

### Construct Design and Cloning

cDNA libraries were generated from total RNA extracts (TRIZol, Invitrogen) from wild-type HH10 chicken embryos, reverse transcribed using Superscript III First-Strand Synthesis (Invtrogen) following manufacturer’s instructions. Full *APOD*, *PAX2*, *ETV4*, and *SOX8* open reading frames were cloned, and XhoI and ClaI restriction enzyme sites were added, by PCR with high-fidelity AccuPrime *Taq* DNA polymerase (Invitrogen). All primer sequences used are presented in Table S1. Overexpression constructs were produced by digestion and ligation into the multiple cloning site of pCI H2B-RFP (Betancur et al., 2011) for *APOD* and pCIG (Megason and McMahon, 2002) for *PAX2, ETV4* and *SOX8*, between XhoI and ClaI sites (all enzymes from New England Biolabs).

### Sequence Analysis

Bulk RNA sequencing analysis was performed by downloading raw read counts from manually dissected chicken otic placodes (GEO accession GSE69185, (Chen et al., 2017)) and imported into R. Duplicate gene identifiers were collapsed by summing counts, lowly expressed genes were filtered out, remaining counts were normalized using TMM factors in edgeR (Robinson et al., 2010), and converted to log2-counts per million (log2-CPM) with a prior count of 1. The resulting log2-CPM values were used for visualization with ggplot2 (Wickham et al.).

FGF-sensitive otic enhancer peaks were explored from previously published H3K27ac ChIP-seq data (Tambalo et al., 2020). Briefly, BED files were downloaded from GEO accession GSE137664, aligned to the chicken genome (galGal4 build) on the UCSC genome browser, and examined at the *APOD* gene locus. The gene structure, control naïve PPR, and FGF-induced tracks were exported and prepared for presentation using Affinity Designer 2 (v2.6.5, Serif).

### Microscopy and Image Analysis

Whole mount and transverse section imaging was performed using a Zeiss AxioImager M2 with Apotome.3 or Zeiss LSM900 Confocal with Airyscan2 Detector. Fluorescent whole mount embryo images were collected and displayed as widefield views or collected as Z-stacks and displayed as maximum intensity projections. Fluorescent transverse section images were displayed as a single optical plane or as maximum intensity projections of Z-stacks. Images were processed, prepared for display, and analyzed using Fiji (Schindelin et al., 2012), and figures were prepared using Affinity Designer (v2.6.5, Serif) or Biorender (Science Suite Inc.). Fluorescence intensity quantitation was performed on maximum intensity projection images in circular ROIs with a diameter of 98.64 µm placed within the center of the otic placode, and otic vesicle areas were measured by drawing a freehand ROI encompassing the vesicle at its largest optical section from Z-stacks. Quantitation of proliferation and apoptosis markers was performed by manually counting PH3- or cleaved caspase 3-positive cells within the otic placode across 2-3 cryosections per embryo and plotting the average results per embryo.

### Statistical Analysis and Rigor

All statistical analysis and plotting were performed using Prism10 (GraphPad), unless otherwise described. For all datasets, normality and lognormality was tested using a Kolmogorov-Smirnov test. Parametric tests (*t* test or one-way ANOVA) were performed to compare normally distributed datasets, and nonparametric tests (Wilcoxon ranks sum) were used for non-normally distributed data. The specific tests are detailed as appropriate in the figure legends. All paired connected points in slope plots represent a single embryo, and every dot in jitter plots represent a single embryo, showing pooled measurements from embryos generated from at least three independent replicates.

