## Supplemental Information for "Lipid-mediated reinforcement of FGF/MAPK signaling enables robust otic placode specification"

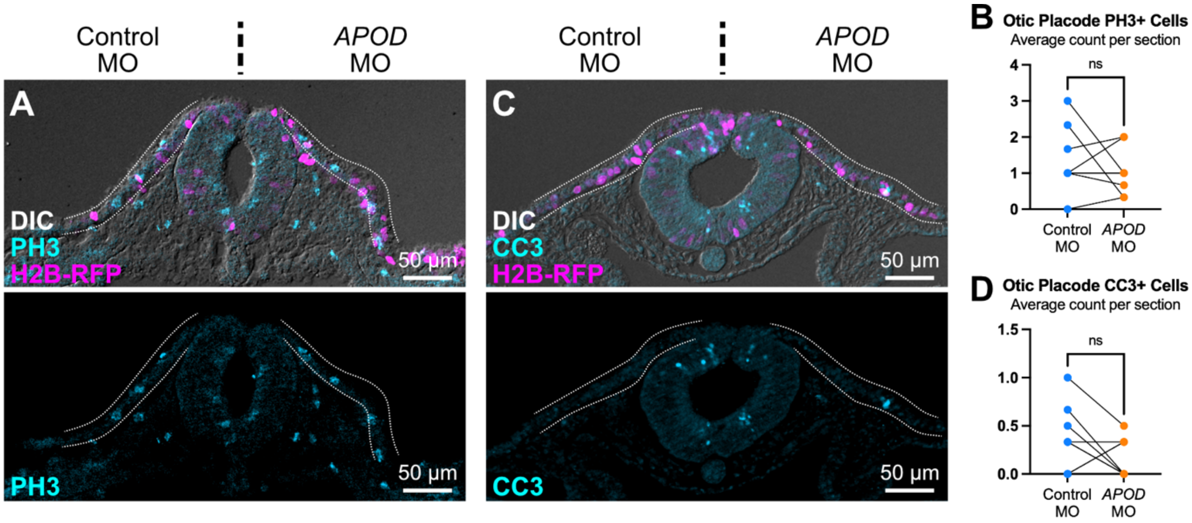

**Figure S1. APOD does not influence proliferation or apoptosis in the otic placode**
**A, C.** Representative transverse sections of *APOD* morphant embryos immunolabeled for phospho-histone H3 (PH3, mitotic cells) and cleaved caspase 3 (CC3, apoptotic cells). **B, D.** PH3- or CC3-positive cells were counted in the otic ectoderm from least three transverse sections per embryo (n=7 embryos for PH3, and n=6 embryos for CC3), and average counts are displayed cell counts per embryo. ns, not significant, Paired t-test.

### Table S1. Key Resources

| REAGENT or RESOURCE | SOURCE | IDENTIFIER |
|---|---|---|
| **Antibodies** | | |
| Rabbit polyclonal anti-SOX10 | Millipore-Sigma | Cat# HPA068898; RRID:AB_2686054 |
| Phospho-p44/42 MAPK (ERK1/2) (Thr202/Tyr204) | Cell Signaling | Cat# 9101; RRID:AB_331646 |
| Rabbit polyclonal anti-phospho-Histone H3 (Ser10) | Millipore | Cat# 06-570, RRID:AB_310177 |
| Rabbit polyclonal anti-Caspase-3, activated form | R & D Systems | Cat# AF835, RRID:AB_2243952 |
| Goat polyclonal anti-GFP | Rockland | Cat# 600-101-215M, RRID:AB_2612804 |
| Rabbit polyclonal anti-RFP/DsRed | MBL International | Cat# PM005, RRID:AB_591279 |
| **Experimental Models: Organisms/Strains** | | |
| Fertilized chicken eggs, White leghorn | UConn Poultry Unit, Storrs, CT | N/A |
| Fertilized chicken eggs, White leghorn | MSU Poultry Teaching & Research Center, Okemos, MI | N/A |
| **Deposited Data** | | |
| Bulk RNA-Seq: Dissected chicken otic placode territory (stages HH6, 5-6ss, 8-9ss, and 11-12ss) | (Chen et al., 2017) | GEO Number: GSE69185 |
| ChIP-Seq: FGF-responsive otic enhancers as called HOMER from genome-wide H3K27ac occupancy (pre-placodal region explants without vs with FGF2 supplementation) | (Tambalo et al., 2020) | GEO Number: GSE137664 |
| **Oligonucleotides** | | |
| Morpholino: Control MO CCTCTTACCTCAGTTACAATTTATA | Gene Tools | N/A |
| Morpholino: APOD MO CACAGCGGCACTTACAGCATCTCCT | Gene Tools | N/A |
| Primer: APOD Kozak FWD XhoI ATACTCGAGGCCACCATGCTGGGCACGGCAGCGCA | This paper | N/A |
| Primer: APOD Stop REV ClaI GCGCATCGATTTACATCTCGGCAGGGCAGTTGAGC | This paper | N/A |
| PRIMER: PAX2 Kozak XhoI FWD ATTGCTCGAGGCCACCATGGATATGCACTGCAAGGC | This paper | N/A |
| Primer: PAX2 Stop ClaI REV ATATATCGATACTAGTGGCGGTCATAGGCAG | This paper | N/A |
| Primer: ETV4 Start XhoI FWD ACGCCTCGAGATGAAGGGGTACGTGGACCAG | This paper | N/A |
| Primer: ETV4 REV Stop ClaI REV ACCTATCGATGCTAGTAGGTGTAGCCTTTGCC | This paper | N/A |
| Primer: SOX8 Kozak XhoI FWD ATTACTCGAGGCCACCATGCTCAACATGACCGAGGAGC | This paper | N/A |
| Primer: SOX8 STOP ClaI REV ATATATCGATTTAAGGCCTCGTCAGGGTTGTG | This paper | N/A |
| **Recombinant DNA** | | |
| Plasmid: pCI::H2B-RFP | (Betancur et al., 2010) | N/A |
| Plasmid: pCIG::nls-GFP | (Megason and McMahon, 2002) | N/A |
| Plasmid: pCI::APOD | This paper | N/A |

| | | |
|---|---|---|
| Plasmid: pCIG::ETV4 | This paper | N/A |
| Plasmid: pCIG::PAX2 | This paper | N/A |
| Plasmid: pCIG::SOX8 | This paper | N/A |
| Plasmid: pCIG::dnFGFR1 | (Stuhlmiller and García-Castro, 2012) | N/A |
| **Software and Algorithms** | | |
| Fiji version 2.16.0/1.54p | (Schindelin et al., 2012) | https://imagej.net/Fiji |
| R version 4.4.3 | (Ihaka and Gentleman, 1996) | https://www.r-project.org/ |
| RStudio version 2023.06.1+524 | (Racine, 2012) | https://www.rstudio.com/ |
| ggplot2 version 4.0.1 | (Wickham et al.) | https://ggplot2.tidyverse.org/ |
| edgeR version 4.4.2 | (Robinson et al., 2010) | https://bioconductor.org/packages/release/bioc/html/edgeR.html |
| GraphPad Prism 10 version 10.6.0 (796) | | https://www.graphpad.com/ |
| ChatGPT (GPT-5) | OpenAI | http://chat.openai.com/ |
